# *Xenopus laevis* as an infection model for human pathogenic bacteria

**DOI:** 10.1101/2025.03.10.642459

**Authors:** Ayano Kuriu, Kazuya Ishikawa, Kohsuke Tsuchiya, Kazuyuki Furuta, Chikara Kaito

## Abstract

Animal infection models are essential for understanding bacterial pathogenicity and corresponding host immune responses. In this study, we investigated whether juvenile *Xenopus laevis* could be used as an infection model for human pathogenic bacteria. *Xenopus* frogs succumbed to intraperitoneal injection containing the human pathogenic bacteria *Staphylococcus aureus*, *Pseudomonas aeruginosa*, and *Listeria monocytogenes*. In contrast, non-pathogenic bacteria *Bacillus subtilis* and *Escherichia coli* did not induce mortality in *Xenopus* frogs. The administration of appropriate antibiotics suppressed mortality caused by *S. aureus* and *P. aeruginosa*. Strains lacking the *agr* locus, *cvfA* (*rny*) gene, or hemolysin genes in *S. aureus*, LIPI-1-deleted mutant of *L. monocytogenes*, which attenuate virulence within mammals, exhibited reduced virulence in *Xenopus* frogs compared to their respective wild-type counterparts. Bacterial distribution analysis revealed that *S. aureus* persisted the blood, liver, heart, and muscles of *Xenopus* frogs until death. These results suggested that intraperitoneal injection of human pathogenic bacteria induces sepsis-like symptoms in *Xenopus* frogs supporting their use as a valuable animal model for evaluating antimicrobial efficacy and identifying virulence genes in various human pathogenic bacteria.

## Introduction

According to a 2019 report, 13.6% of global deaths were associated with infections caused by bacterial pathogens such as *Staphylococcus aureus* and *Pseudomonas aeruginosa* (1). *S. aureus* is a Gram-positive pathogenic bacterium that resides in the nasal cavity of approximately 30% of healthy adults and causes pneumonia, osteomyelitis, and endocarditis (1). Drug-resistant *S. aureus*, known as methicillin resistant *S. aureus* (MRSA), is spreading worldwide, necessitating the development of new treatments (2). *P. aeruginosa* is a bacterium commonly found in environments such as water and soil and in the intestinal tract of healthy humans. However, during infection, it can cause urinary tract infections, sepsis, and other diseases. Most antibiotics are ineffective against infections caused by multidrug-resistant *P. aeruginosa* (3). *Listeria monocytogenes* is found in river water and animal intestinal tracts and causes meningitis, septicemia, and fetal septic granuloma in foodborne infections (4). In the United States, approximately 20% of all foodborne deaths are attributed to *L. monocytogenes* infections (5). To establish treatment strategies for these bacterial infections, understanding bacterial pathogenic mechanisms and host immune system to counter the bacterial infection is crucial.

Mammalian animal models such as mice have traditionally been used to elucidate bacterial and host factors involved in the bacterial infection process. These models have the advantage of having immune systems similar to those of humans; however, ethical and high-cost issues limit the use of large numbers. Recently, a global movement has been initiated to reduce animal testing and use alternative methods, as evidenced by the 2013 EU ban on the sale of cosmetics tested on animals (6). Alternatives to animal testing, including *in vitro* and *in silico* experiments and use of cultured mammalian cells, have attracted attention (7, 8); however, they fail to reproduce the complex interactions occurring in animal bodies. Animal infection models remain necessary to investigate bacterial-host interactions and disease progression leading to death.

To avoid ethical and cost problems associated with mammals, invertebrates, such as nematodes (9), fruit flies (10), silkworms (11), honey worms (12), and crickets (13), as well as vertebrate fish like zebrafish (14, 15), are being used as infection models (16, 17). These animals allow large-scale testing of bacterial pathogenicity and antimicrobial agents. However, invertebrates lack antibody-mediated adaptive immune mechanisms, making it impossible to analyze bacterial interactions with the adaptive immune system. In addition, their open vasculature, unlike vertebrates, uniquely affects the pathways for bacterial spread *via* body fluids. While zebrafish possess immune systems relatively similar to those of humans (14, 15), their respiratory system, which relies on gill breathing, differs significantly from that of humans.

Establishing a non-mammalian infection model with new advantages is expected to advance research on infectious diseases. *Xenopus laevis*, an amphibian vertebrate, has been widely used in developmental biology for several decades (18). *X. laevis* harbors organs, organ arrangements, and circulatory systems similar to those of humans (19). Moreover, its immune system closely resembles that of humans and possesses key factors of innate and acquired immunity, including macrophages, neutrophils, B cells, T cells, immunoglobulins, complement system, and antimicrobial peptides (20, 21). These characteristics suggest that *X. laevis* could be useful for analyzing bacterial dissemination and immune interaction comparable to those in humans. The entire genome sequence of *X. laevis* has been completed (22), and genome editing using TALEN and CRISPR Cas9 is now possible (23–25). *Xenopus* frog can be kept at a high densities in aquariums without aeration because of their lung breathing (26). Fifty juvenile *Xenopus* frogs, each approximately 3 cm in length, can be maintained in about 10 L of water. In addition, they can be easily injected using human clinical tuberculin syringes without anesthetic treatment, as ice cold anesthesia is effective. These features provide marked advantages in experiments that require large number of animals to test using injections. *Xenopus* frogs have been used as an infection model for *Mycobacterium marinum* (19), which infects various aquatic organisms. However, they have not yet been employed as infection models for human pathogenic bacteria. Furthermore, no reports have demonstrated the use of *Xenopus* frogs to evaluate bacterial virulence genes or antimicrobial agent efficacy. In this study, we aimed to establish a *Xenopus* frog infection model for human pathogenic bacteria.

Animal infection models should be effective for evaluating bacterial virulence genes. The *agr* locus (27) and *srtA* (28) and *cvfA* (*rny*) genes (29) encode important *S. aureus* virulence factors. The *agr* locus encodes quorum sensing-related proteins and regulates toxin and cell surface protein expression (30). The *srtA* gene is required to anchor cell wall proteins involved in bacterial adhesion to host cells (31). The *cvfA* gene encodes RNase Y, which is essential for hemolysin production (29, 32, 33). Similarly, Listeria Pathogenicity Island 1 (LIPI-1) region in *L. monocytogenes* (34) encodes *prfA*, *plcA*, *hly*, *mpl*, *actA*, and *plcB* (35, 36). Hemolysin LLO (*hly*) (37) and phospholipase C (PLC) (*plcA* and *plcB*) (38) facilitate *L. monocytogenes* escape from phagosomes or autophagy (35, 39). These virulence genes of *S. aureus* and *L. monocytogenes* are crucial for bacterial pathogenicity in animal models such as mice (40–43), rabbits (44), crickets (13), silkworms (28, 42), and nematodes (45, 46). Therefore, these bacterial genes are suitable for examining the effectiveness of novel animal infection models to evaluate bacterial virulence. Animal infection models should also be effective for assessing the efficacy of antimicrobial agents. In this study, methicillin-susceptible and methicillin-resistant *S. aureus* strains, and drug-susceptible and multidrug-resistant *P. aeruginosa* strains were used to evaluate the efficacy of clinically used antimicrobial agents.

## Results

### Intraperitoneal injection of *S. aureus* can kill *Xenopus* frogs

To determine whether *Xenopus* frogs are susceptible to human pathogenic bacteria, bacterial suspensions of *S. aureus* were intraperitoneally injected into the *Xenopus* frogs. The frogs died within 24 h post-injection and their entire body exhibited reddening (**Fig. 1A**).

**Fig. 1.**
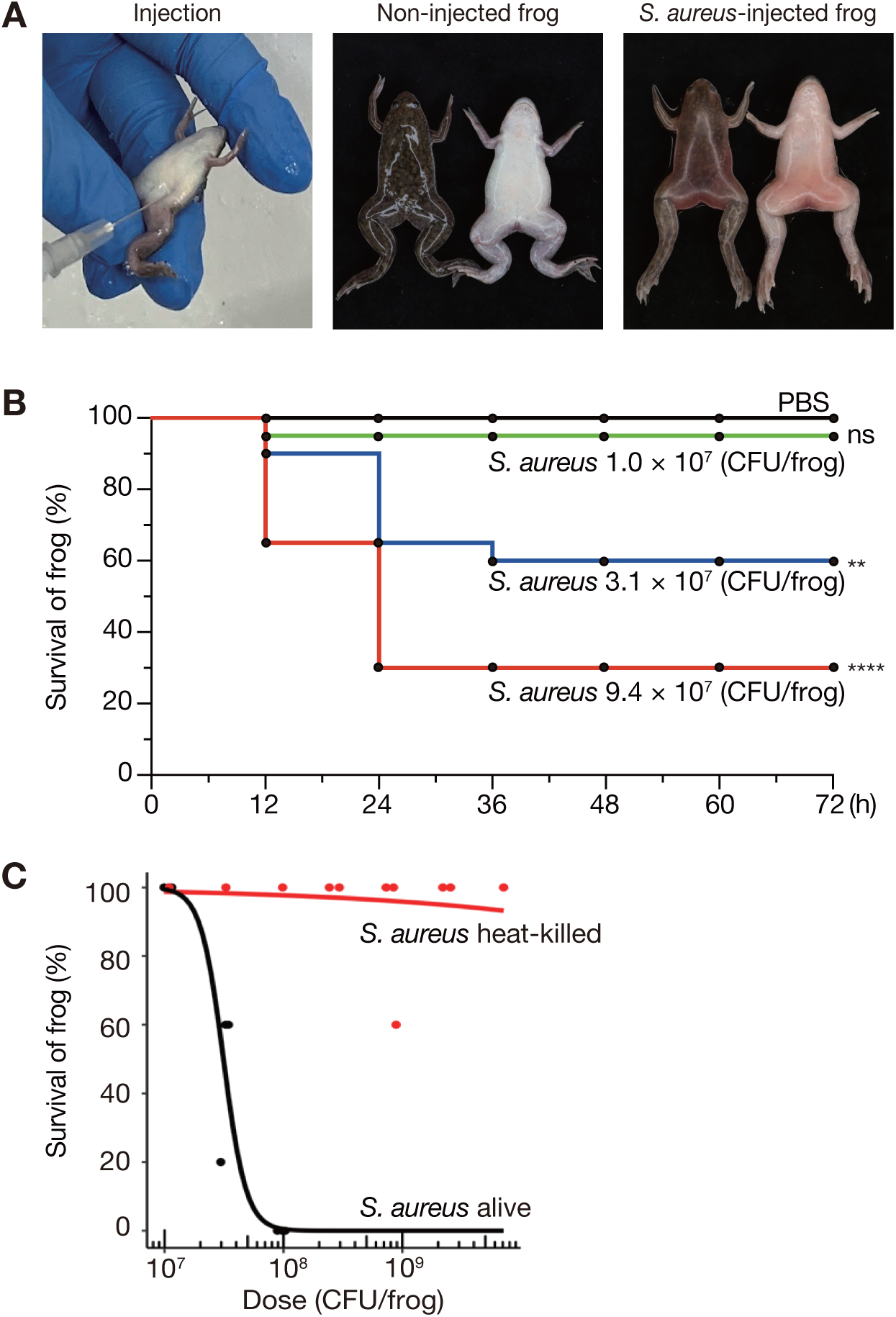
Intraperitoneal injection of *S. aureus* is lethal to *Xenopus* frogs. (A) Intraperitoneal injection of frogs is shown (left panel). Frogs injected with *S. aureus* NCTC8325-4 strain (2.8 × 10^8^ CFU/frog) were dead within 24 h post-injection (right panel). Frogs injected with PBS survived (center panel). (B) Time-course survival analysis of frogs intraperitoneally injected with varying doses of *S. aureus* NCTC8325-4 strain. Five frogs were injected per sample, and survival data were pooled from four independent experiments (n=20). ns: p > 0.05, **: p < 0.01, ****: p < 0.0001. (C) Dose-response survival curve of frogs injected with live or heat-killed *S. aureus* NCTC8325-4. Overnight culture of *S. aureus* was autoclaved at 121°C for 20 min before use as heat-killed bacteria. Five frogs were injected per sample and LD_50_ was determined from survival curve pooled from three independent experiment data.

To assess whether mortality was dependent on the number of injected bacteria, frogs were injected with varying concentrations of *S. aureus* suspension, and their survival rate was observed over time. Mortality was observed between 12 and 36 h after injection, which increased with higher bacterial concentrations (**Fig. 1B**). These results indicated that *S. aureus* infection leads to frog death in a dose-dependent manner.

To confirm that the death of frogs post-injection was caused by live bacteria, we examined whether heat-killed *S. aureus* exhibited similar lethal activity. The survival rate of frogs injected with live *S. aureus* decreased depending on the number of bacteria at 24 h post-injection, whereas that of frogs injected with heat-killed *S. aureus* did not decrease at 24 h post-injection (**Fig. 1C**). The median lethal dose (LD_50_) of live and heat-killed *S. aureus* were 3.2 × 10^7^ and 7.1 × 10^9^ CFU/frog, respectively. These results suggested that frogs died owing to the biological activity of viable *S. aureus*, such as bacterial growth and toxin production.

### Various human pathogenic bacteria can kill *Xenopus* frogs

To determine whether the frog-killing ability is conserved among human pathogenic bacteria other than *S. aureus* NCTC8325-4, we examined the virulence of other *S. aureus* strains, *P. aeruginosa*, and *L. monocytogenes* against *Xenopus* frogs. Intraperitoneal injection of MRSA (MRSA8), *P. aeruginosa* strain PAO1, multidrug-resistant *P. aeruginosa* (BAA-2114), and *L. monocytogenes* strain EGD caused frog death based on the bacterial number (**Fig. 2A, 2B, 2C, 2D**), similar to *S. aureus* NCTC8325-4 (**Fig. 1C**). Additionally, we examined whether laboratory strains of non-pathogenic bacteria such as *Escherichia coli* (47) and *Bacillus subtilis* (48) induces mortality. Intraperitoneal injections of *E. coli* BW25113 and *B. subtilis* 168 *trpC2* showed little killing activity to frog within the tested bacterial concentrations (**Fig. 2E, 2F**). LD_50_ calculated from survival curves showcased that *P. aeruginosa* BAA-2114 had the highest LD_50_ of 1.8 × 10^9^ CFU/frog, whereas *S. aureus* NCTC8325-4 and *L. monocytogenes* EGD exhibited lower LD_50_ values (< 4.3 × 10^7^ CFU/frog) (**Fig. 2G**). LD_50_ values of *E. coli* BW25113 and *B. subtilis* 168 *trpC2* were not determined, as they exceeded 3.3 × 10^9^ and 1.6 × 10^9^ CFU/frog, respectively (**Fig. 2G**). These results suggested that nonpathogenic bacteria have limited lethality in frogs, whereas human pathogenic bacteria demonstrate frog-killing ability, which varies among bacterial species and strains.

**Fig. 2.**
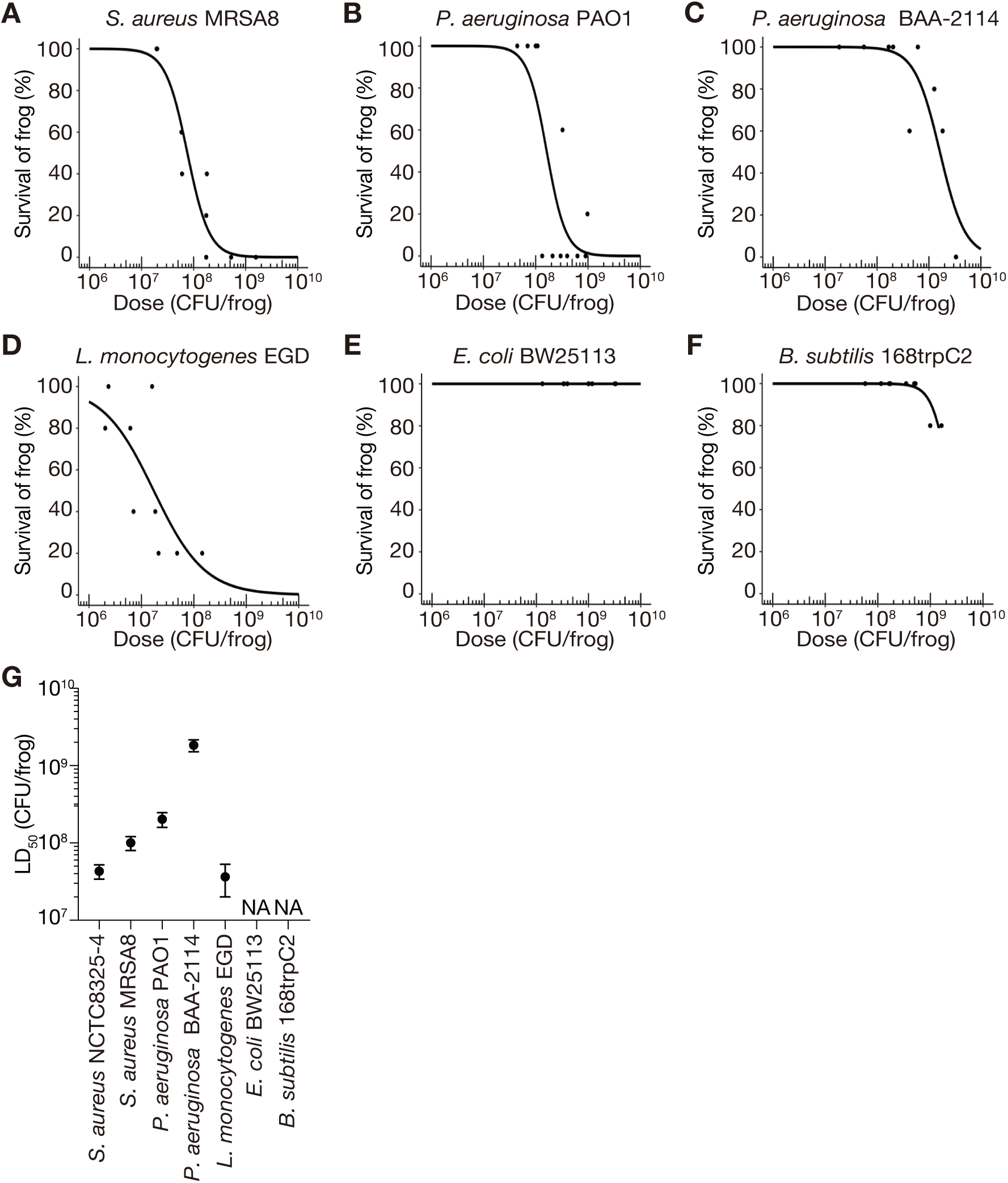
Various species of human pathogenic bacteria induce mortality in *Xenopus* frogs. (A–F) Dose-response survival curves of frogs injected intraperitoneally with *S. aureus* MRSA8 (A), *P. aeruginosa* PAO1 (B), *P. aeruginosa* BAA-2114 (C), *L. monocytogenes* EGD (D), *E. coli* BW25113 (E), and *B. subtilis* 168trpC2 (F) generated from the survival rate at 24 h post-injection. Five frogs were injected with three different concentrations of bacterial suspensions in a single experiment, and survival data obtained from three independent experiments are shown. (G) Based on the survival data (A–F), logistic regression analysis was performed using the statistical software R to determine LD_50_. Error bars indicate standard error. LD_50_ for NCTC8325-4 was calculated from the survival data in Fig. 1C.

### Evaluation of antibacterial drug efficacy in *Xenopus*

To determine whether *Xenopus* frogs can serve as a model for evaluating antimicrobial efficacy, we examined the therapeutic effects of antimicrobials on frog death induced by human pathogenic bacteria. First, the toxicities of kanamycin (KM), oxacillin (OX), vancomycin (VCM), ciprofloxacin (CPFX), and ceftazidime (CAZ) were tested, which confirmed that none of the tested antibiotics caused frog mortality at the antimicrobial concentrations used in this study (**Table S1**). Antimicrobials were administered immediately after bacterial injection, and frogs were considered recovered if their survival rate was greater than 50% at 120 h post-infection.

MRSA8, an MRSA strain of *S. aureus*, is resistant to oxacillin (β-lactam) and kanamycin (aminoglycoside), but remains sensitive to vancomycin (11, 13, 49). BAA-2114 is a multidrug-resistant *P. aeruginosa* isolated from the sputum samples of patients with cystic fibrosis (50), which displays resistance to kanamycin, ceftazidime (β-lactam), and ciprofloxacin (fluoroquinolones) (54–56).

Frogs injected with NCTC8325-4, a methicillin-susceptible strain of *S. aureus*, without antibacterial injections exhibited a survival rate of less than 10%, whereas administration of KM, OX, or VCM after injecting bacterial suspension resulted in survival rates of > 50% at 120 h post-infection (**Fig. 3A**). Frogs injected with *S. aureus* MRSA8 had 0% survival at 120 h post-infection, whereas those injected with VCM after MRSA8 achieved a 100% survival rate in the same time (**Fig. 3B**). In contrast, injecting KM or OX following MRSA8 resulted in nearly 0% survival at 120 h post-infection (**Fig. 3B**). These results indicated that NCTC8325-4 infection in frogs can be treated with KM, OX, and VCM, whereas MRSA8 infection respond only to VCM.

**Fig. 3.**
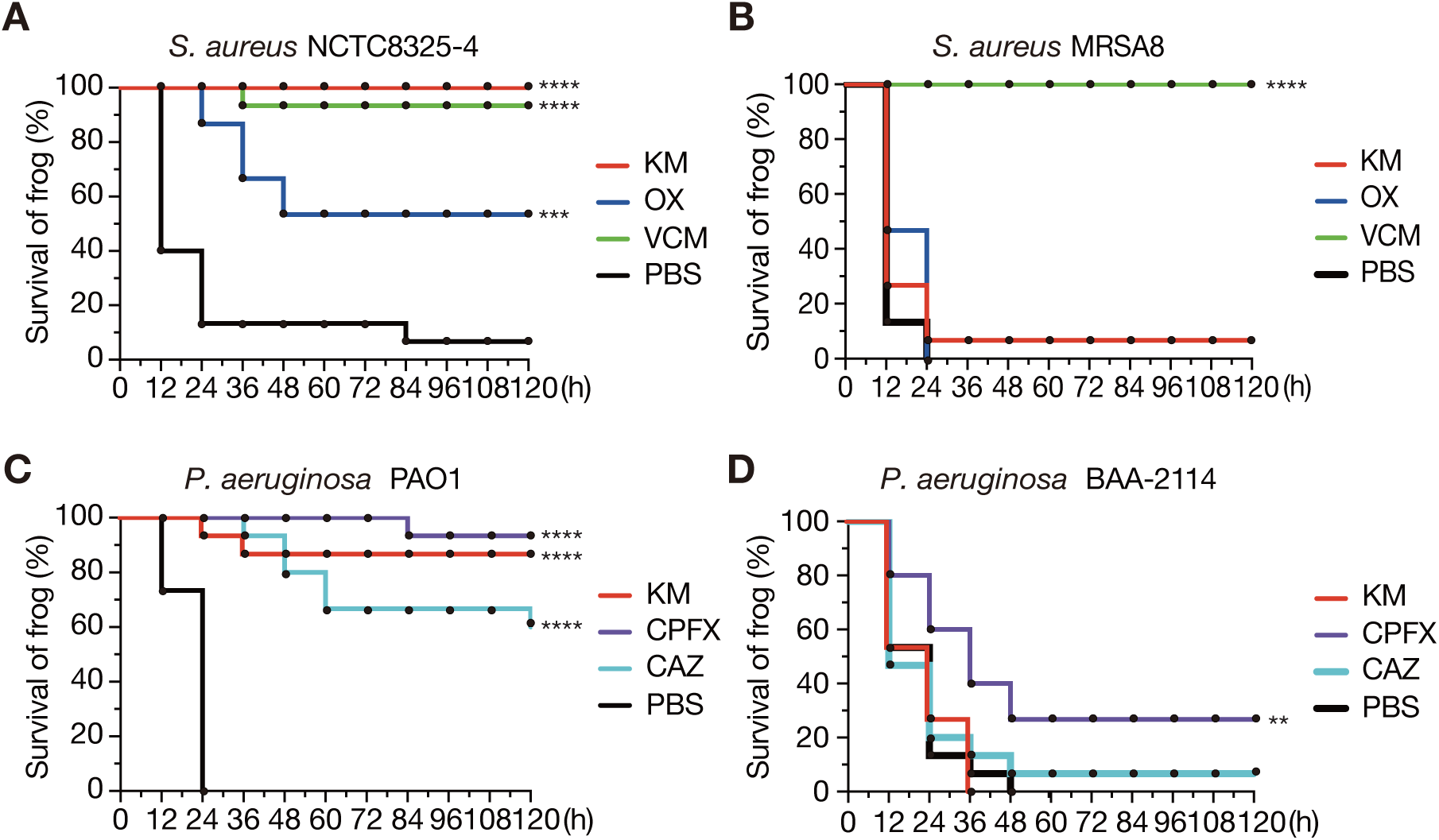
Antimicrobial agents inhibit bacterial infections in *Xenopus* frogs. Therapeutic effects of kanamycin (KM), oxacillin (OX), and vancomycin (VCM) on frogs injected with *S. aureus* NCTC8325-4 (A) and MRSA8 (B) were examined. Similarly, the effects of kanamycin (KM), ciprofloxacin (CPFX), and ceftazidime (CAZ) on frogs infected with *P. aeruginosa* PAO1 (C) and BAA-2114 (D) were examined. Immediately after intraperitoneal injection of bacterial suspensions, frogs were intraperitoneally administered with 50 µL of 2 mg/mL antimicrobial agent (100 µg/frog) or PBS, and survival was measured every 12 h. Five frogs were used per agent, and survival data were pooled from three independent experiments (n=15). Significant differences between PBS-- and antimicrobial-injected groups were determined using log-rank test (* p < 0.05, ** p < 0.01, *** p < 0.001, **** p < 0.0001).

Frogs injected with *P. aeruginosa* PAO1 without antimicrobials showed 0% survival at 120 h post-infection, while those treated with KM, CPFX, or CAZ following PAO1 administration, showed survival rates of > 50% (**Fig. 3C**). Similarly, frogs injected with *P. aeruginosa* BAA-2114 without antimicrobial agents was 0% at 120 h post infection, whereas those injected with KM, CPFX, or CAZ after BAA-2114 showed less than 50% survival rates in the same timeframe (**Fig. 3D**). These findings suggested that PAO1 infections in frogs can be effectively treated with KM, CPFX, or CAZ, whereas BAA-2114 infections are refractory to these antimicrobial agents.

### Evaluation of bacterial virulence genes

To determine whether the virulence genes of pathogenic bacteria could be evaluated using *Xenopus* frogs, virulence gene deletion mutants of *S. aureus* and *L. monocytogenes* were intraperitoneally injected into *Xenopus* frogs, and their viability was measured 24 h post-infection. *S. aureus* NCTC8325-4 mutants with deleted *agr* locus (27), *srtA* (28), and *cvfA* (29) killed frogs in a dose-dependent manner (**Fig. 4A**). The LD_50_ values of *agr*- and *cvfA*-deleted mutants were more than 10-fold higher than those of the wild-type strain (**Fig. 4A**). In contrast, the LD_50_ of *srtA*-deleted mutant was nearly the same as that of the wild-type strain (**Fig. 4A**). These results suggested that *agr* and *cvfA* contribute to *S. aureus* virulence against *Xenopus* frogs, while *srtA* does not.

**Fig. 4.**
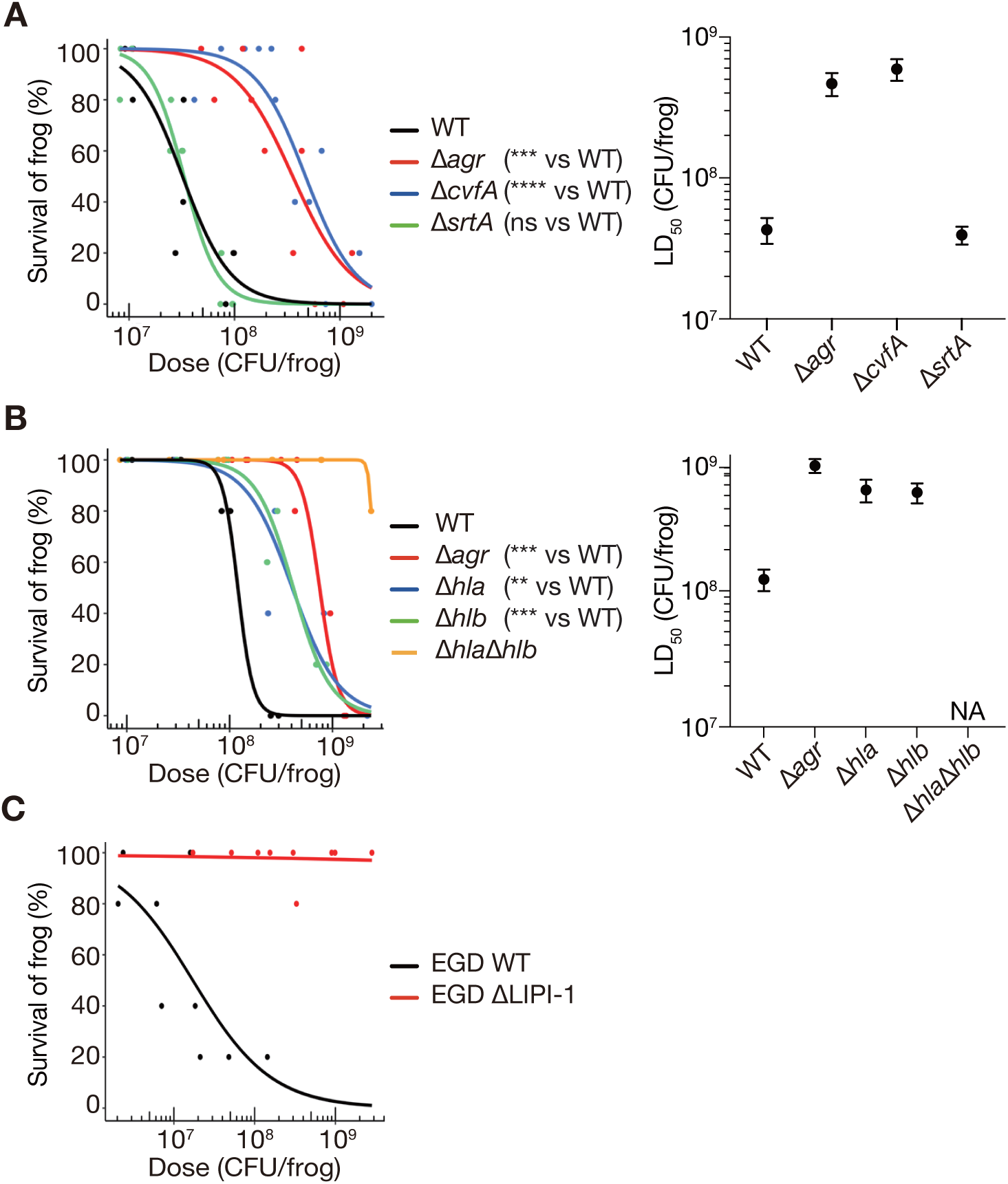
Deletion of bacterial virulence genes reduces bacterial killing activity against *Xenopus* frogs. (A) Comparison of the frog-killing abilities of *S. aureus* NCTC8325-4 strain (WT) and strains with deleted *agr*, *srtA*, and *cvfA*. (B) Comparison of the frog-killing abilities of NCTC8325-4 WT and strains with deleted *agr*, *hla*, *hlb*, and *hla*/*hlb*. (C) Comparison of the frog-killing ability of *L. monocytogenes* EGD (WT) and a strain with deleted of LIPI-1. The data for *L. monocytogenes* EGD WT are the same as those presented in Fig. 2A. For all panels, bacterial suspensions were injected intraperitoneally into frogs, and survival was measured at 24 h post-injection. Five frogs were injected with each of three different concentrations of bacterial suspensions per strain in a single experiment. Dose-response survival curves were generated using logistic regression based on data from three independent experiments, and LD50 and standard error were calculated. Significant differences between gene-deleted and WT strains were determined using the likelihood ratio test (**: p < 0.01, ***: p < 0.001, ****: p < 0.0001). Error bars represent standard error.

Given that the *agr-* and *cvfA*-deleted mutants attenuated virulence against *Xenopus* frogs and *agr* and *cvfA* are required for hemolysin production (30, 57), we hypothesized that the reduced hemolysin production in these mutants may contribute to their attenuated virulence. To verify this, we assessed the virulence of hemolysin gene deletion mutants in *Xenopus* frogs. Deletion mutants of *hla* (encoding α-hemolysin) and *hlb* (encoding β-hemolysin) are known to reduce virulence in mammals (28, 44). The LD_50_ values of *hla*- and *hlb*-deleted mutants were higher than those of the wild-type strain (**Fig. 4B**). In addition, the LD_50_ of the *hla*/*hlb* double deletion mutant exceeded 2.4 × 10^9^ CFU/frog, which was higher than the LD_50_ values of mutants harboring either deletion (**Fig. 4B**). These results suggested that *hla* and *hlb* act in an additive manner to kill *Xenopus* frogs.

*Xenopus* frogs injected with the LIPI-1-deleted mutant of *L. monocytogenes* EGD showed minimal mortality at 24 h post-infection (**Fig. 4C**). The LD_50_ of wild-type strain was 3.6 × 10^7^ CFU/frog, whereas that of the LIPI-1-deleted mutant was greater than 2.8 × 10^9^ CFU/frog **(Fig. 2G, 4C**). These results suggested that the LIPI-1 locus of *L. monocytogenes* is essential for its virulence against *Xenopus* frogs.

### Intraperitoneal injection of *S. aureus* into *Xenopus* frogs causes bacterial dissemination

We examined the visceral morphology of *Xenopus* frogs intraperitoneally injected with *S. aureus* NCTC8325-4 and compared it with that of non-injected frogs (**Fig. 5A**). The stomachs of non-injected frogs were pale pink, and the muscles were soft and movable. In contrast, the stomachs of injected frogs, examined immediately after death at 7 h post-injection, were red and deformed, and rigor mortis had set in, with hard and immobile muscles (**Fig. 5A**). In addition, blood obtained by amputating the upper legs of non-injected frogs was dark red, whereas that of frogs immediately after death appeared light red.

**Fig. 5.**
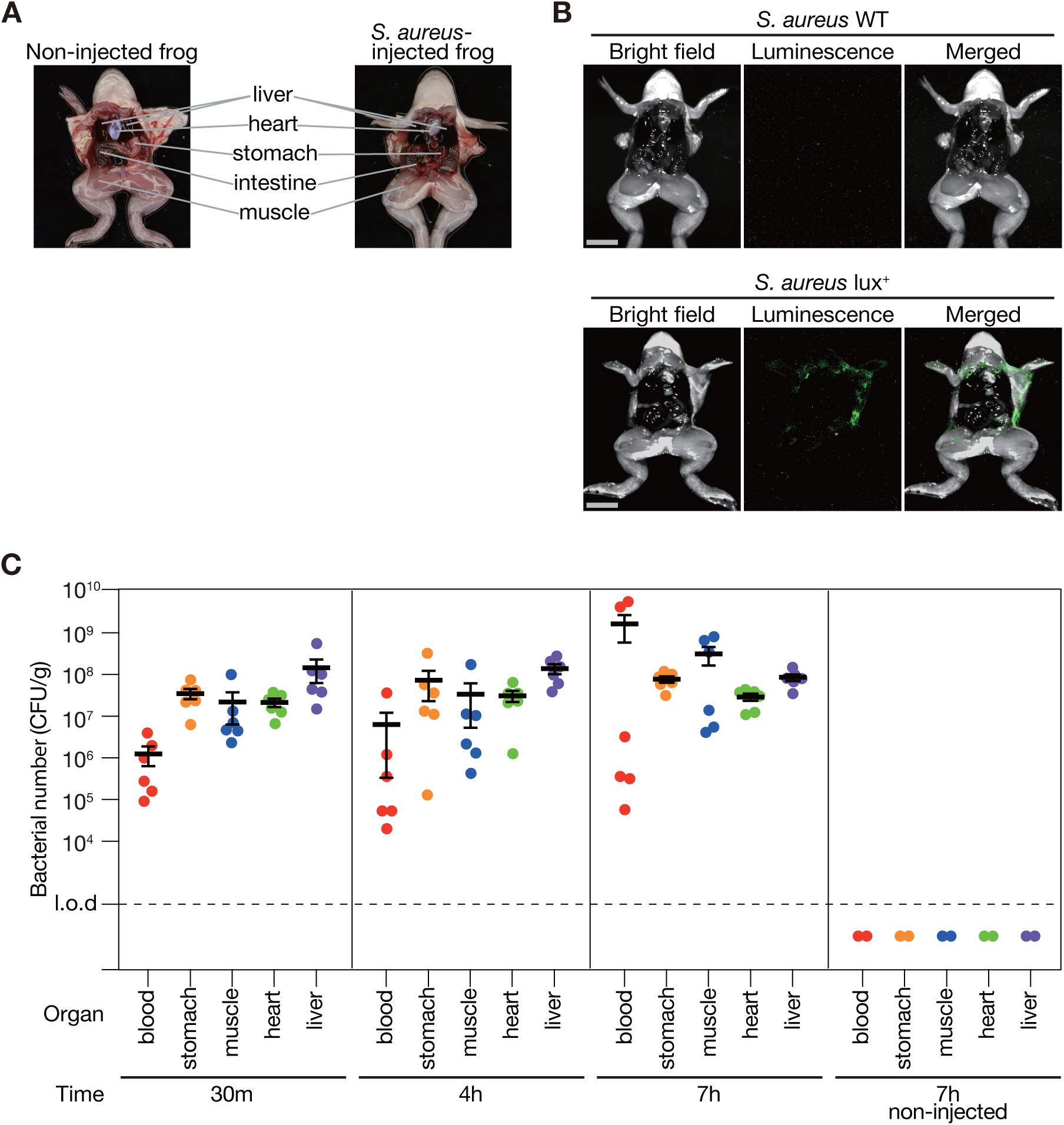
Bacteria disseminate from intraperitoneal cavity to whole body in *Xenopus* frogs. (A) Photograph of a *Xenopus* frog with its abdomen open. The left shows a non-injected frog. The right shows a frog that died 7 h after intraperitoneal injection of *S. aureus* NCTC8325-4 strain (8.6 × 10^8^ CFU/frog). (B) *S. aureus* NCTC8325-4 strain (WT) or NCTC8325-4 strain carrying pND50-gmkp1-lux was injected intraperitoneally into frog, and the abdomen was opened 7 h post-injection to observe luminescence. Bright field (left), luminescence (center), and merged images (right) are displayed. Scale bar: 10 mm. The bacterial dose administered was 8.6 × 10^8^ CFU/frog. (C) *Xenopus* frogs were intraperitoneally injected with *S. aureus* NCTC8325-4 strain (8.6 × 10^8^ CFU/frog), and bacterial CFUs were quantified in blood and organs at 30 min, 4 h, and 7 h post-injection. At each time point, three animals were analyzed in a single experiment, and two independent experiments were performed. Mean CFU count from six frogs at each time point is represented by a solid horizontal bar, with the error bar indicating standard error. Open symbols indicate data from PBS-injected control frogs (n=2).

To investigate the distribution of *S. aureus* in *Xenopus* frogs after intraperitoneal injection, frogs were intraperitoneally injected with *an S. aureus* strain expressing the *lux* operon, which enables a luminescence reaction without the need for external substrates. Seven h post-infection, luminescence was observed at the periphery of the abdominal cavity (**Fig. 5B**). Conversely, no luminescence was detected in frogs administered with *S. aureus* strain lacking the *lux* operon (**Fig. 5B**). These results suggested that *S. aureus* intraperitoneally injected into *Xenopus* frogs disseminated throughout the abdominal cavity.

To further investigate the systemic distribution of *S. aureus* in *Xenopus*, *Xenopus* frogs were intraperitoneally injected with *S. aureus* and dissected at 30 min, 4 h, and 7 h post-injection to quantify bacterial counts in the blood and various organs (stomach, leg muscle, heart, and liver). *S. aureus* were detected at all time points and in all examined organs, with bacterial numbers maintained from 30 min to 7 h post-infection (**Fig. 5C**). In contrast, *S. aureus* were not detected in the organs of the frogs injected with PBS (**Fig. 5C**). Therefore, all detected *S. aureus* were derived from the intraperitoneally injected bacteria. Notably, *S. aureus* were observed in the blood, and in the leg muscles that does not directly contact the abdominal cavity, suggesting that intraperitoneally injected *S. aureus* spread throughout the body *via* blood.

## Discussion

This study demonstrated that *Xenopus* frogs are susceptible to intraperitoneal injection of the human pathogenic bacteria, *S. aureus*, *P. aeruginosa*, and *L. monocytogenes* and lead to mortality. Furthermore, clinically effective antimicrobial agents suppressed frog death infected with *S. aureus* and *P. aeruginosa*, while bacterial mutant strains with deleted virulence genes in *S. aureus* and *L. monocytogenes* exhibited attenuated virulence. This study is the first to suggest that *Xenopus* frogs are a viable animal infection model for evaluating both the efficacy of antimicrobial agents and virulence gene function of human pathogenic bacteria.

This study showed that *S. aureus* disseminated into the blood and organs of *Xenopus* frogs following intraperitoneal injection, resulting in systemic infections. In humans, intraperitoneal infections caused by laparotomy or intestinal obstruction can progress to sepsis (58, 59). In addition, intraperitoneally injected bacteria have been detected in the blood and liver of mouse infection model (60, 61). These observations suggested that the intraperitoneal bacterial infection in *Xenopus* frogs can simulate bacterial sepsis in mammals, where intraperitoneally infected bacteria disseminate throughout the body.

In this study, we found that *hla*, *hlb*, as well as *agr* and *cvfA*, which regulate the transcriptional expression of *hla* and *hlb*, are required for the lethality of *S. aureus* in *Xenopus* frogs (**Fig. 4A, 4B**). *S. aureus* strains lacking *hla* and *hlb* showcase reduced virulence in rabbits (44). In human alveolar epithelial cells, α-hemolysin interacts with the host cell surface protein ADAM10 and forms a heptameric β-barrel structure, creating pores in the cell membrane (62). ADAM10 knockout in mouse alveolar epithelial cells confers resistance to lethal pneumonia caused by *S. aureus* (60). β-Hemolysin is a neutral sphingomyelinase that degrades sphingomyelin in cell membranes (64). According to Xenbase, a database of *X. laevis* and *X. tropicalis*, *X. laevis* has ADAM10 (target of α-hemolysin) and a sphingomyelin synthase (target of β-hemolysin), suggesting that α-hemolysin and β-hemolysin may exert similar effects in *X. laevis* as in mammals. These findings support the use of *Xenopus* frogs for analyzing bacterial virulence mechanisms mediated by α- and β-hemolysins.

During intraperitoneal infection in *Xenopus* frogs, the *srtA*-deleted mutant in this study showed virulence comparable to that of the wild-type *S. aureus* (**Fig. 4A**). In rats and mice, the pathogenicity of *srtA*-deleted mutant is lower than that of the wild-type strain administered via various methods, including intraperitoneal injection (62–64). Similarly, in silkworms and crickets, the *srtA*-deleted mutant is less virulent than the wild-type strain in systemic infection models induced by intrahemolymph injection (13, 28). In contrast, in the nematode infection model, where bacteria were fed to the nematodes, *srtA*-deleted mutant displayed similar pathogenicity to that of the wild-type strain (68, 69). As *srtA* deficiency attenuates virulence in mice following intraperitoneal injection (65), this suggests that the non-attenuated virulence of *srtA*-deleted mutant observed in *Xenopus* frogs is not owing to the infection route, but rather to host animal-specific differences. The *srtA* gene is involved in anchoring more than 10 different cell wall adhesion proteins (70, 71). The molecular mechanisms underlying why cell adhesion by these cell wall proteins is not required for intraperitoneal infection in *Xenopus* frogs need to be analyzed in the future.

The mortality of *Xenopus* frogs caused by drug-susceptible and drug-resistant strains of *S. aureus* and drug-susceptible strains of *P. aeruginosa* was suppressed by the administration of appropriate antimicrobial agents. However, multidrug resistant *P. aeruginosa* remained lethal despite antimicrobial treatment (**Fig. 3**). Hence, *Xenopus* frogs could prove useful for evaluating antibiotic efficacy against sepsis caused by *S. aureus* and *P. aeruginosa*. *Xenopus* juvenile frogs are smaller than mice, which enables the evaluation and exploration of antimicrobial agents using small drug doses. In this study, antimicrobials were intraperitoneally injected immediately after injecting bacterial suspensions. Future studies should evaluate the efficacy of other antimicrobial administration routes used in clinical practice, such as oral and subcutaneous administration. The *Xenopus* infection model has potential for identifying and analyzing key human pathogenic bacterial genes involved in sepsis progression and evaluating therapeutic interventions.

## Materials and Methods

### Ethics statement

This study was conducted in strict accordance with the Fundamental Guidelines for Proper Conduct of Animal Experiments and Related Activities in Academic Research Institutions under the jurisdiction of the Ministry of Education, Culture, Sports, Science and Technology, 2006. All mouse protocols complied with the Regulations for Animal Care and Use of Okayama University and were approved by the Animal Use Committee at Okayama University (approval number: OKU-2024557).

#### Xenopus laevis

*X. laevis* juvenile frogs were purchased from a breeding company (Xenopus Yoshoku Kyozai, Ibaraki, Japan). The frogs were 3–4 cm in size and weighed approximately 2.5 g. Fifty frogs were housed in a 13.6 L (37 cm W × 22 cm D × 25 cm H) aquarium, which was half-filled with water. The aquarium was maintained at a room temperature of 22°C. The frogs were fed with dried goldfish food (Cat. 4971453050347, Itosui, Tokyo, Japan) every two days. The rearing water was replaced with fresh dechlorinated water once every two days (72).

### Bacterial strains

All bacterial strains used in this study are listed in Table 1. *S. aureus* gene deletion mutants for *agr* locus (73), *srtA* (28), *cvfA* (29), *hla* (28), *hlb* (28), and *hla*/*hlb* (28) were constructed in our previous studies. We confirmed the deletion of the respective genes by PCR, using genomic DNA as the template and oligonucleotide primers (**Table 2**). *L. monocytogenes* mutant strain lacking LIPI-1 was constructed according to a previously described method (74). The downstream regions of *prfA* and *plcB* were amplified via PCR and ligated into pHS-MCS. *L. monocytogenes* EGD strain was electroporated with the obtained plasmids, resulting in a LIPI-1-deleted mutant. The genomic DNA of *L. monocytogenes* was extracted using the QIAamp Blood Mini Kit (QIAGEN,Venlo, Netherlands) as previously described (75), and LIPI-1 deletion was confirmed through PCR.

**Table 1.**
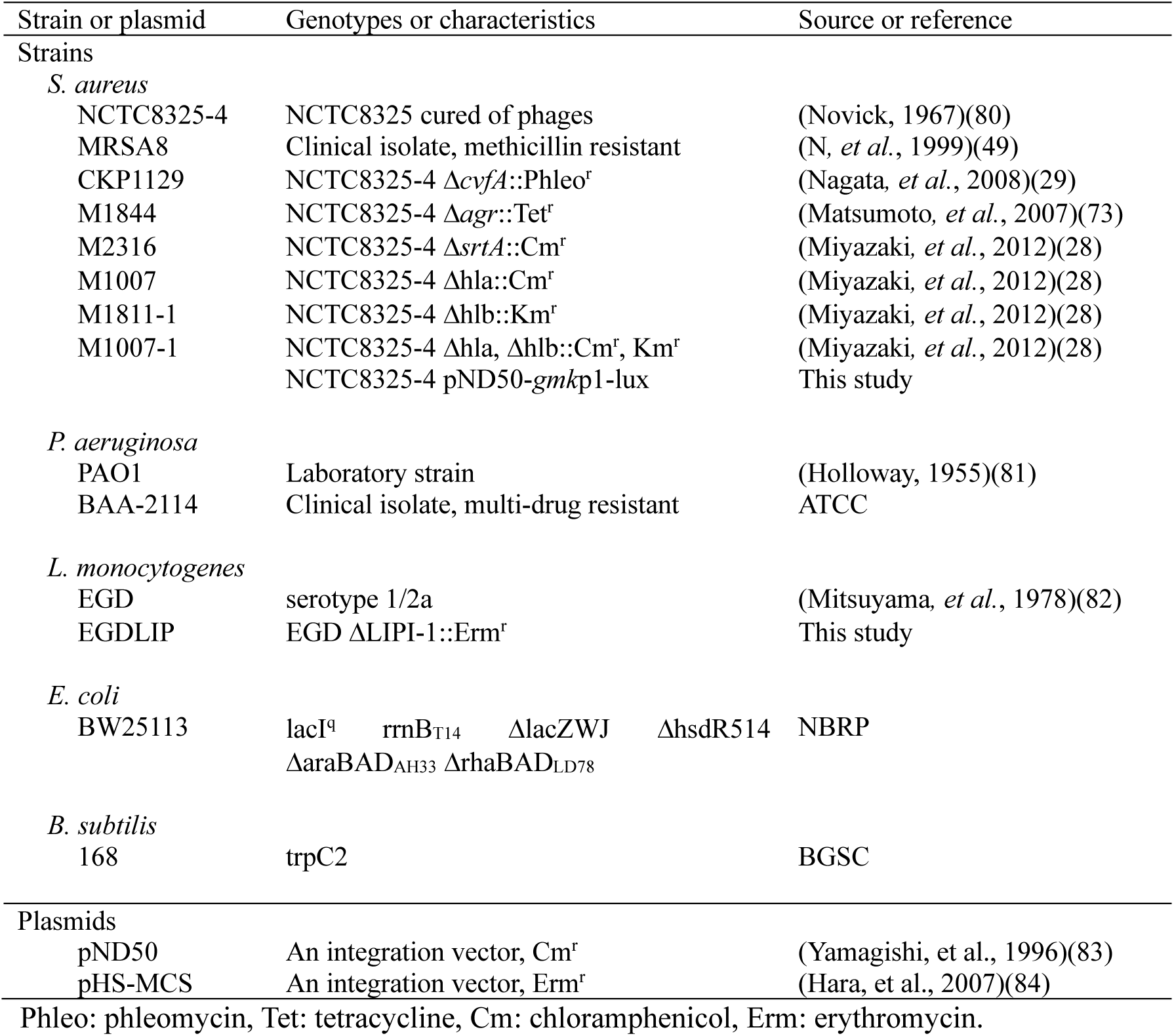
List of the bacterial strains and plasmids used.

**Table 2.**
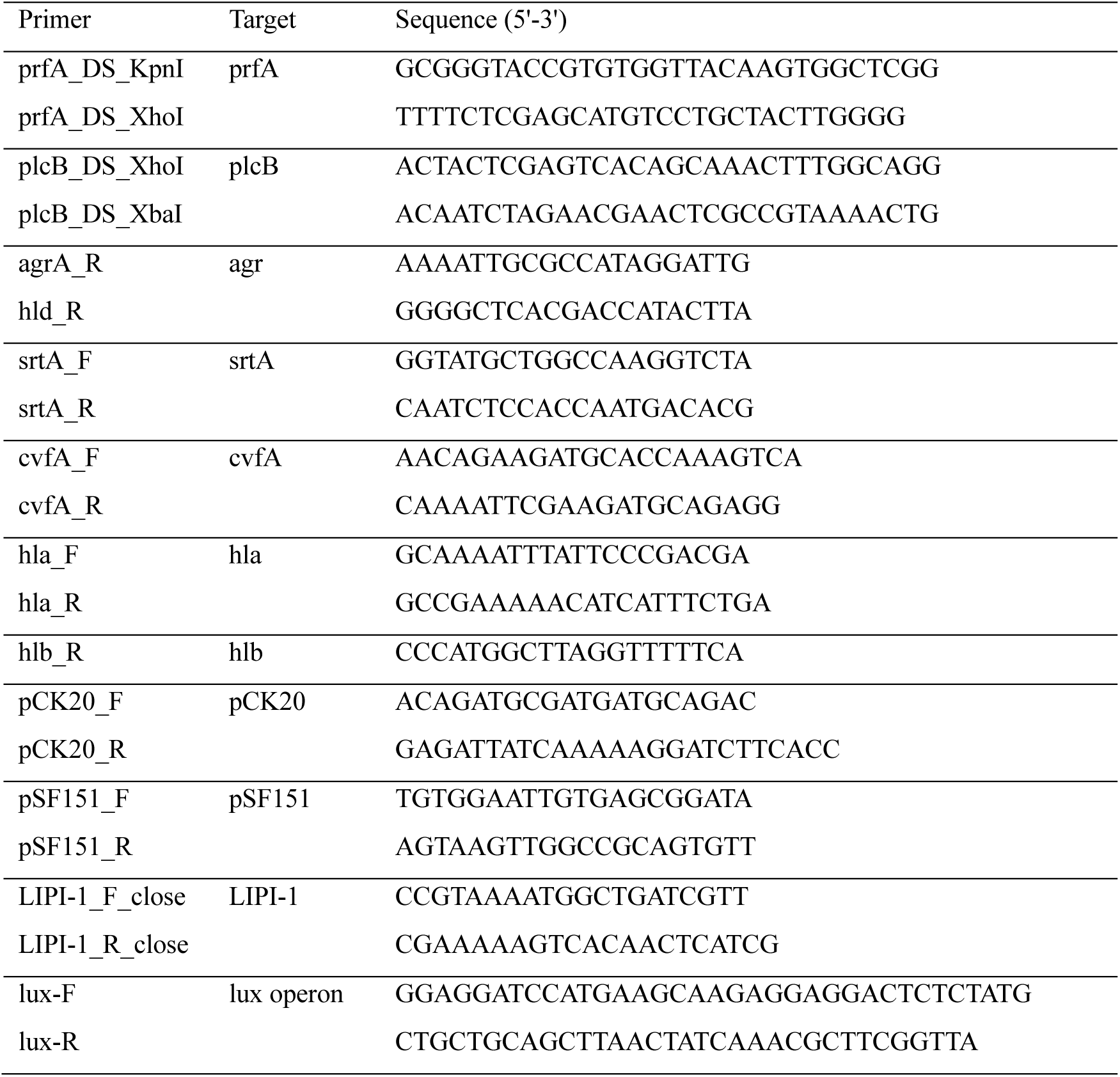
Primers used in this study.

### Preparation of bacterial solution

A single *S. aureus* colony on tryptic soy agar was inoculated into 5 mL of tryptic soy broth medium and aerobically incubated at 37°C overnight. The overnight culture was centrifuged at 10,400 × *g* for 10 min, and the bacterial pellet was resuspended in phosphate-buffered saline (PBS) and diluted to prepare bacterial solutions at appropriate concentrations. The same procedure was performed for *P. aeruginosa* and *L. monocytogenes*, using Lysogeny broth and brain heart infusion media, respectively. In multiple independent experiments, the optical density at 600 nm (OD_600_) of overnight cultures was measured, and a calibration curve was generated to determine the relationship between CFU and OD_600_. This calibration curve was used to estimate the CFU injected into frogs based on the OD_600_ of overnight cultures in the infection experiments.

### Preparation of antimicrobial solutions

Antimicrobial solutions were prepared as follows: 10 mg each of kanamycin monosulfate (TCI, Tokyo, Japan), oxacillin (SIGMA, St. Louis, MO), vancomycin hydrochloride (FUJIFILM, Tokyo, Japan), and ceftazidime hydrate (LKT, St. Paul, MN) were dissolved in 5 mL PBS to achieve a concentration of 2 mg mL^−1^. Ciprofloxacin hydrochloride monohydrate (TCI, Tokyo, Japan) was dissolved in Milli-Q water to a concentration of 2 mg mL^−1^. All antimicrobial solutions were stored at -30°C and thawed before use.

### Intraperitoneal injections

*Xenopus* frogs were anesthetized by placing them on crushed ice for 2 min. A 50 µL bacterial solution was injected into the abdominal cavity of *X. laevis*, just above the left leg, using a 27G needle (outer diameter) of 3/4-inch length (Terumo, Tokyo, Japan) (**Fig. 1A**) attached to a 1 mL tuberculin syringe (Terumo, Tokyo, Japan). Five animals per sample were injected, placed in a 1 L beaker containing 500 mL water, and kept at 27°C in the dark. The frogs recovered from cold anesthesia and begin swimming immediately after being placed in the beaker. To exclude the possibility of death due to low-temperature anesthesia or injection-related stress, a negative control group injected with PBS was prepared for all experiments.

### Calculation of LD_50_

Three different concentrations of bacterial solutions were employed for each experiment. Overnight cultures were centrifuged at 10,400 × *g* for 10 min, the bacterial pellet was resuspended in PBS, and a three-fold dilution series was prepared using PBS. Five animals were injected per sample, and survival rates were recorded at 24 h post-infection. The experiment was performed thrice, and the survival rate data and injected CFU were plotted on a graph. A dose-response survival curve was created using logistic regression using the statistical software R to obtain LD_50_ and standard error. Survival data for each experiment are presented in **Table S1**.

### Luminescence measurement

DNA fragments containing *lux* operon were amplified by PCR using genomic DNA from *S. aureus* Xen29 (76) as template and a primer pair (**Table 2**). The amplified DNA was inserted into the BamHI and PstI recognition sites of a plasmid carrying the *gmk* promoter region (77), resulting in pND50-*gmk*p1-lux. *S. aureus* RN4220 strain was transformed with pND50-*gmk*p1-lux through electroporation, and the plasmid was subsequently transferred into NCTC8325-4 strain using phage 80α via transduction. The NCTC8325-4 strain carrying pND50-*gmk*p1-lux was intraperitoneally injected into *Xenopus* frogs. At 7 h post-injection, the abdomen was opened and chemiluminescence was measured using a CCD camera (iBright 1500, Thermo Fisher, Waltham, MA). The camera was configured to an exposure time of 10 min, zoom 3.0, and focus 390. Three images were captured per animal under the same conditions, and the average function of ImageJ was used to compute the mean luminescence intensity from the three photographs.

### Measurement of bacterial count in organs

*Xenopus* frogs were intraperitoneally injected with *S. aureus* NCTC8325-4. At 30 min, 4 h, and 7 h post-injection, the frogs were anesthetized by immersing them in 2 g L^−1^ ethyl m-aminobenzoate methanesulfonate solution (Nacalai Tesque, Kyoto, Japan) for 5 min. The right upper leg was amputated, and blood was collected. Muscles of the right upper leg, stomach, heart, and liver were excised. To remove residual blood, the organs other than blood were washed with PBS, weighed, and then homogenized in 400 µL PBS using TissueLyser II (QIAGEN) at 25 Hz for 1.5 min. A 10-fold dilution series of blood and organ homogenates were prepared using 0.9% NaCl, Approximately 5 µL of each dilution was spread onto mannitol salt agar plates, a selective medium for *S. aureus*, and incubated at 37°C overnight. The colonies were counted according to a previously defined method (78). Bacterial numbers per gram of blood and organ were calculated based on colony counts and dilution ratios.

### Statistical Analysis

The log-rank (Mantel-Cox) test was performed using GraphPad Prism software (ver. 10.2.3) to determine significant differences in the time-survival curves. A dose-response survival curve was generated via logistic regression to plot frog survival against CFU injected into frogs using R for Windows (ver. 4.3.3). The LD_50_ and standard error were estimated using the glm function from the MASS library (79). A likelihood ratio test was conducted using the anova function to determine significant differences between the two dose-response survival curves. The R code used for these analyses is registered in the GitHub repository (URL: https://doi.org/10.5281/zenodo.14978657, doi: 10.5281/zenodo.14978657).

## Acknowledgements

This study was supported by the Japan Society for the Promotion of Science (JSPS) Grants-in-Aid for Scientific Research (Grants 22K14892, 23K24131, 23K06130, 24K01760, 24K21872). We thank the National BioResource Project-E. coli (National Institute of Genetics, Japan) for providing the *E. coli* BW25113 strain and BGSC (Bacillus Genetic Stock Center) for providing *B. subtilis* 168 strain.

